# Orally Delivered dsRNA-Derived siRNAs Reach the Central Nervous System in *Leptinotarsa decemlineata*

**DOI:** 10.64898/2026.03.11.711085

**Authors:** Venkata Partha Sarathi Amineni, Doga Cedden

**Affiliations:** Institute of Phytomedicine, Department of Applied Entomology, University of Hohenheim, Otto-Sander-Straße 5, 70599 Stuttgart, Baden-Württemberg, Germany; Branch Bioresources, Fraunhofer Institute for Molecular Biology and Applied Ecology IME, Giessen, Germany

**Keywords:** RNA interference (RNAi), double-stranded RNA (dsRNA), Argonaute 2 (AGO2), systemic RNAi, central nervous system (CNS), blood–brain barrier (BBB), Leptinotarsa decemlineata

## Abstract

RNA interference (RNAi) has emerged as an eco-friendly approach to pest management and relies on the processing of exogenous double-stranded RNA (dsRNA). RNAi-based pest management is highly effective in the Colorado potato beetle (*Leptinotarsa decemlineata*); however, the tissue-specific distribution and processing of exogenous dsRNA following oral uptake remain incompletely understood. In this study, we investigated whether ingested dsRNA reaches the central nervous system (CNS) and is processed into active small interfering RNAs (siRNAs). Adult beetles were fed dsmGFP-coated leaf disks, and RISC-bound small RNAs were isolated from midgut, CNS, and remaining body tissues using a RISC-enrichment approach. Small RNA sequencing revealed abundant 21-nucleotide antisense guide-strand siRNAs in all analysed tissues, with relative proportions following the order midgut > CNS > remaining tissues. Notably, antisense siRNAs of consistent size were detected in CNS samples, indicating that exogenous dsRNA or its processed products can access neural tissue and enter the RNAi silencing machinery. These findings provide strong biochemical evidence that orally taken-up dsRNA is processed into AGO-loaded siRNAs in the *L. decemlineata* CNS. Together, our results offer a tissue-resolved view of functional RNAi activity in this species and contribute to a mechanistic understanding of systemic dsRNA transport in coleopteran pests.

**Graphical abstract:** 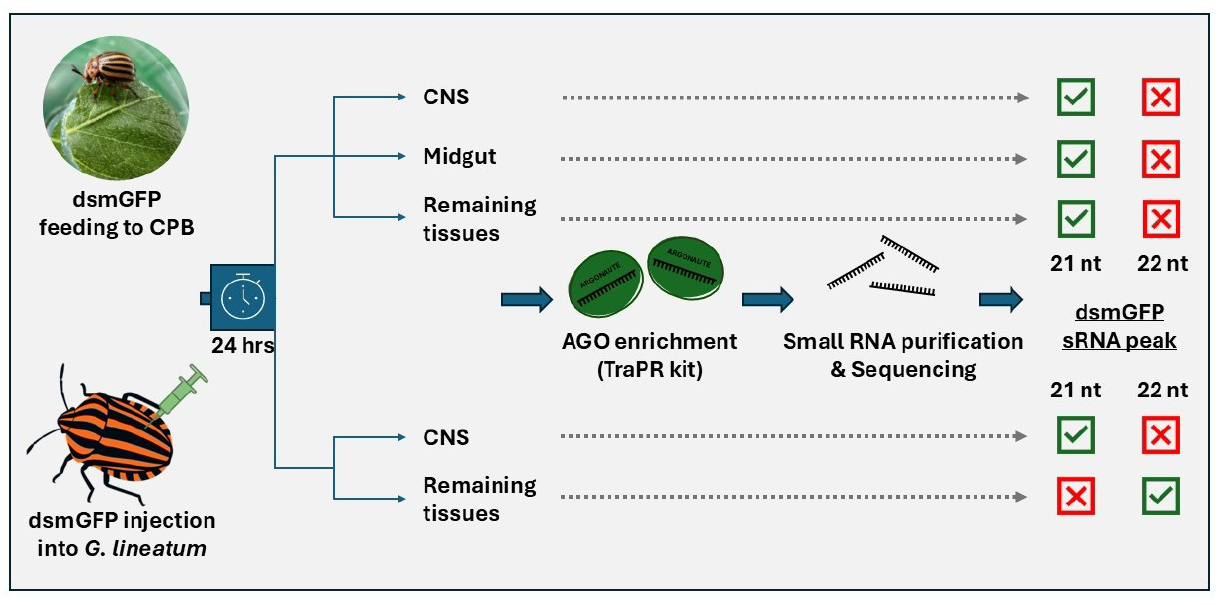

## Introduction

RNA interference (RNAi) has emerged as a powerful molecular tool for post-transcriptional gene silencing and is an emerging pest control strategy with tremendous potential for agricultural applications^1,2^. RNAi is initiated by processing of double-stranded (ds)RNA by the Ribonuclease III enzyme Dicer-2 into mainly 21-24 nucleotide long double-stranded small interfering (si)RNAs in the cytoplasm of the cell^3^. These siRNAs are later incorporated into Argonaute, thereby forming the RNA-induced silencing complex (RISC)^4^, where the guide siRNA strand will remain and serve to find the target mRNA transcript for AGO-2 mediated cleavage^5^. RNAi-based pest control techniques can be very precise to target pest species and can be designed to have a lower risk to non-target organisms using bioinformatics^6,7^,unlike many conventional chemical insecticides with broad range of target species^8^.

Colorado potato beetle (*Leptinotarsa decemlineata*, Coleoptera: Chrysomelidae) is one of the highly sensitive insect pests for foliar applied dsRNA with very high RNAi based gene silencing efficacy at relatively low dosages^9–11^. It served as an ideal model organism to study the RNAi mechanism exhibiting robust systemic RNAi responses following both injection and oral delivery of dsRNA^10^. Previous studies have showed that orally delivered dsRNA can induce effective knockdown of several lethal target genes including the genes encoding *vATPase subunit A*^12^, *Proteasome* subunits^10^, *Actin*^13^, *Mesh*^14^ and several others^11^ causing significant mortalities and development abnormalities. However, despite these findings, several fundamental knowledge gaps exist in spatial distribution, tissue specific processing and particularly the neural penetration of ingested dsRNA and its derived siRNAs.

To induce a systemic RNAi response in insect, the foliar applied dsRNA must be successfully taken up by the epithelium of intestine after oral ingestion, followed by the transport further to other distant organs from the primary site of uptake. In previous studies, multiple dsRNA uptake mechanisms were proposed in insects, including SID-like channel proteins in CPB^15^, and Clathrin mediated endocytosis^12^ through scavenger receptors on the cell membranes^16^. Post-oral uptake of dsRNA, it is still unclear how exactly is the dsRNA is spreading systemically throughout the insect. A critical and not well studied question is whether the orally delivered dsRNA can penetrate the blood-brain barrier to reach the central nervous system (CNS) in sufficient quantities to establish functional RNAi in neural tissues. In the desert locust (*Schistocerca gregaria*), the injection of double-stranded RNA (dsRNA) led to strong gene silencing in brain tissue, whereas a very low level of target gene silencing was observed in the reproductive organs^17^. However, direct evidence demonstrating the entry, processing of dsRNA into central nervous system (CNS) of other insects and target neural gene silencing following the oral dsRNA delivery is very limited.

In insects, the blood-brain barrier (BBB) is an important physiological barrier that separates the central nervous system from the haemolymph^18,19^. The insect BBB maintains ionic homeostasis critical for neuronal function, primarily via septate-junction–mediated restriction of paracellular diffusion and regulated transport across barrier glia^18–20^. In addition, ABC efflux transporters expressed at the BBB contribute to chemical defence by exporting many xenobiotics and toxins, limiting their accumulation in the CNS^21–23^. Previous research on plant toxins (cardenolide)-exposed insects has demonstrated that despite high haemolymph toxin concentrations, nervous tissue can remain protected because the BBB, formed by the perineurial/subperineurial glial sheath, strongly restricts toxin entry, via a diffusion barrier for polar cardenolides (e.g., ouabain) and active efflux for more lipophilic cardenolides (e.g., digoxin)^23^. This highlights the BBB’s role as a robust defensive barrier that can strongly limit CNS penetration of toxins such as cardenolides. The question of whether orally delivered exogenous dsRNA can access neural tissue by traversing these barrier mechanisms or enter the CNS through alternative pathways such as receptor-mediated endocytosis or other dsRNA transport mechanisms, remains unclear.

Investigating whether exogenously delivered dsRNA or its derived siRNAs reach neural tissues and successfully interacting with core RNAi machinery in the CNS will have substantial implications for both understanding systemic RNAi biology and optimizing RNAi-based pest control strategies. Most studies assessing exogenous dsRNA processing rely on whole-body^10^ or gut tissue^12,24^ total small RNA sequencing, potentially masking the tissue-specific differences in siRNA accumulation and Dicer processing. To address these knowledge gaps, we used the TraPR RISC-enrichment method (Lexogen GmbH, Vienna, Austria), a validated method for isolating Argonaute (AGO)-bound active small RNAs^25^, combined with next-generation small-RNA sequencing techniques to provide direct biochemical proof of RISC-loaded final active siRNAs in multiple tissues of the CPB following oral delivery of non-target dsRNA-GFP.

Our study focused on determining whether (a) ingested dsRNA is transported through tissue barriers to access the CNS, and (b) AGO-protein/RISC-loaded siRNAs accumulate in CNS tissue with tissue-specific siRNA length and proportion distribution patterns. Using an AGO-enriched small RNA sequencing technique, we found that orally fed dsRNA-derived small RNAs are enriched in CNS tissue, demonstrating that dsRNA can enter the CNS. Extending beyond coleopterans, we also detected enriched dsRNA-derived small RNAs in a hemipteran insect, *Graphasoma lineatum*, following haemolymph injection of dsRNA. This mechanistic evidence establishes a framework for understanding systemic dsRNA trafficking and neural tissue accessibility in coleopteran insect pests, with direct applications for improving RNAi-based pest management strategies.

## Results & Discussion

### dsRNA-GFP-derived siRNA length distribution across tissues

Across all RISC-enriched libraries, sequencing yielded a total of between 1.1 and 21.8 million reads per sample on average, providing robust coverage for small RNA profiling (see Table S 2). Of these reads, 3,600–159,873 mapped to the dsmGFP sequence, corresponding to 500–16,779 reads per million (RPM). In most samples, 21-nt siRNAs were the most abundant size class (43.9–55.6% of mapped reads), with 22-nt siRNAs representing the second most abundant population (11.1–20.8%) (see Table S 2).

The midgut exhibited the highest abundance of 21-nt siRNAs (>4,000 reads per million [RPM]), consistent with this tissue being the primary site of dsRNA uptake and processing. Lower levels were detected in the remaining tissues (∼300–500 RPM) and CNS (∼200–400 RPM), although the 21-nt peak was consistently present. In addition, a minor peak at 34 nt was observed in remaining tissues and midgut samples, in line with previous studies in CPB, whereas this peak was largely absent from CNS samples (Fig. 1).

**Figure 1.**
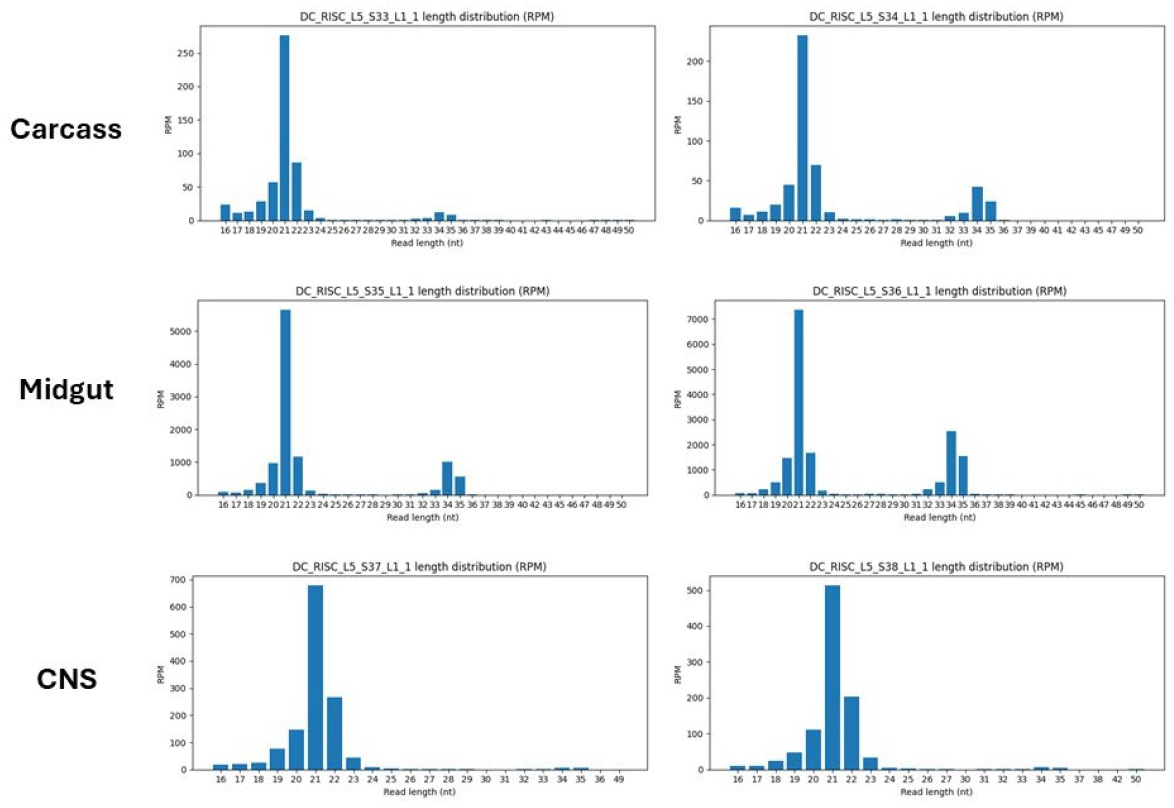
Length distribution of siRNAs in distinct tissue compartments of Colorado Potato Beetle. Bar graphs display read counts (RPM, reads per million) for siRNAs of varying nucleotide lengths in remaining tissues, midgut, and central nervous system (CNS) tissue extracts. Each tissue sample is represented by two biological replicates (left and right panels). Across all tissues and replicates, siRNAs predominantly peak at 21 nucleotides, indicating conserved Dicer processing. Quantitative differences in siRNA abundance are observed, but the position of the peak is indicative of conserved processing preferences, suggesting highly similar dsRNA-processing specificity in each compartment. These results support the conclusion that canonical siRNA generation mechanisms operate highly similar throughout peripheral and neural tissues.

The predominance of 21-nt siRNAs across tissues suggests that dsmGFP is processed through the conserved eukaryotic RNA interference pathway. Similar size profiles have been reported in CPB neonates^10^ and gut tissues^12,24^ following dsmGFP exposure, as well as in other coleopteran species, including *Tribolium castaneum* and *Psylliodes chrysocephala*^26^.

#### Systemic distribution and CNS accessibility of dsRNA-derived siRNAs

RISC-bound siRNAs were detected in all analysed tissues, including the CNS (Fig. 1), indicating that orally administered dsRNA, or its processed intermediates, reaches neural compartments. The quantitative distribution pattern (midgut > remaining tissues > CNS) reflects the primary sites of dsRNA uptake and subsequent systemic transport.

The high abundance of siRNAs in the midgut is consistent with initial exposure at the intestinal epithelium, where dsRNA is internalised through endocytic and/or transporter-mediated mechanisms^15^. The presence of measurable siRNA levels in the CNS, despite anatomical and physiological barriers, demonstrates that dsRNA-derived molecules are distributed systemically rather than being restricted to primary recipient tissues. This finding provides molecular evidence that neural tissues are susceptible to orally taken up dsRNA.

#### Strand-specific mapping of 21-nt siRNAs along the dsmGFP sequence

To identify Dicer cleavage patterns, 21-nt siRNA reads were mapped to the dsmGFP sequence in all tissue types (Fig. 2). Both sense and antisense siRNAs were generated from distinct dsRNA regions rather than being uniformly distributed along the template. Importantly, the positions of these regions were conserved across tissues and biological replicates, despite substantial differences in overall siRNA abundance.

**Figure 2.**
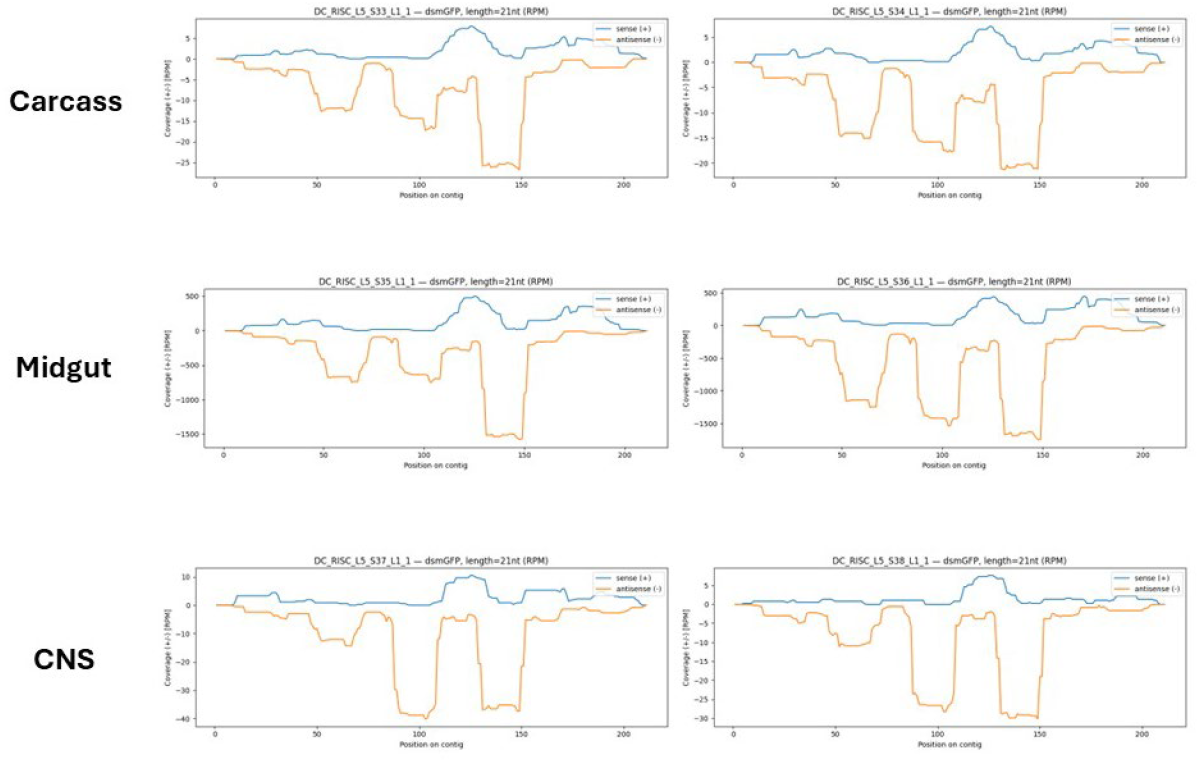
Positional distribution of 21-nt siRNAs mapping to the dsmGFP sequence across tissue compartments. Line plots display normalized read coverage (reads per million, RPM) of 21-nucleotide siRNAs at each position along the dsmGFP sequence in remaining tissues, midgut, and central nervous system (CNS), with two biological replicates per tissue. Curves above the baseline (positive values) indicate read depth for the sense (+) strand, while curves below the baseline (negative values) indicate read depth for the antisense (–) strand. Negative values on the y-axis do not signify negative read numbers but are a convention to represent antisense strand coverage for clarity. All compartments show conserved siRNA hotspot positions with similar strand profiles between tissues and replicates, reflecting highly similar Dicer specificity. Quantitative differences reflect siRNA abundance rather than differences in cleavage sites.

Three major antisense hotspots were detected at approximately 47–67 bp, 85–110 bp, and 130–150 bp along the dsmGFP sequence. The primary hotspot (130–150 bp) represented the most abundant cleavage site in all tissues. These regions closely correspond to hotspots previously identified in studies using the same dsmGFP construct in *T. castaneum* and *P. chrysocephala*^26^.

The conservation of these cleavage sites indicates that Dicer-2-mediated processing operates with defined sequence- and structure-dependent specificity that is largely independent of tissue context. Although RPM values varied among tissues, the dsRNA positions of generated siRNAs remained stable (Fig. 2).

Structural studies have shown that Dicer cleaves dsRNA mainly in regular 21-nt intervals in insects while recognising sequence motifs and local secondary structures that influence cleavage site selection^26,27^. The presence of discrete hotspots in the dsmGFP suggests that terminal structures, sequence composition, and local RNA folding patterns collectively determine preferred processing sites. Furthermore, differences in the half-lives of individual siRNAs derived from the same dsRNA may also contribute to these hotspots.

#### Sense strand configuration and RISC strand selection

Mapping of sense-strand-derived siRNAs revealed a major hotspot at approximately 115–135 bp, which overlapped with or was adjacent to the dominant antisense hotspot. However, sense-strand read depths were consistently lower than antisense reads in all tissues (Fig. 2).

The conserved positioning of sense-strand hotspots across tissues further indicates that Dicer cleavage patterns are tissue-independent, whereas strand bias is introduced primarily at the level of RISC loading or retention.

#### Tissue-specific siRNA accumulation

Quantification of dsmGFP-derived siRNAs revealed pronounced tissue-dependent accumulation (Fig. 3). The midgut contained the highest proportion of dsmGFP-derived sequences (mean = 0.0136), followed by the CNS (mean = 0.0011) and remaining tissues (mean = 0.0005). This distribution reflects the primary role of the midgut in dsRNA uptake and the subsequent dissemination of processed products to peripheral and neural tissues.

**Figure 3.**
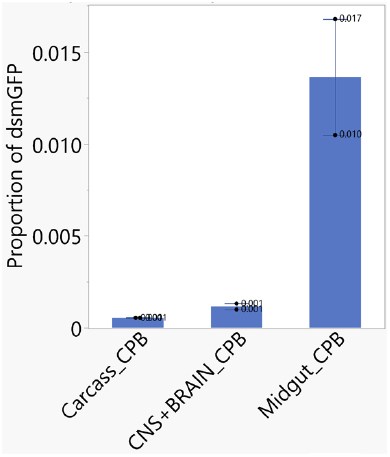
Comparative abundance of dsmGFP-derived siRNAs across tissue compartments. Bar graph displays the proportion of 21-nucleotide siRNAs mapped to the dsmGFP sequence relative to total sequencing reads (reads per million, RPM) in three tissue types: remaining tissues, central nervous system (CNS + BRAIN), and midgut. Data represent mean proportions with error bars indicating standard error of the mean (SEM) from two biological replicates. The midgut shows the highest dsmGFP-derived siRNA proportion, while the remaining tissues and CNS + BRAIN show substantially lower proportions. These results indicate tissue-dependent accumulation of orally delivered dsRNA, with the midgut as the primary site of processing and the CNS demonstrating detectable systemic penetration despite anatomical barriers.

Because dsmGFP-derived siRNAs accounted for ≤1.3% of the total small RNA pool even under high-dose exposure, endogenous small RNAs clearly predominated. These results indicate that the RNAi machinery was not saturated under the experimental conditions. This observation is consistent with previous biosafety studies, which suggest that pathway saturation occurs only at extremely high experimental doses and is unlikely under realistic exposure scenarios. Furthermore, this is consistent with the fact that endogenous small RNAs are usually produced by Dicer-1 and bound by AGO-1, whereas exogenous dsRNAs are processed by Dicer-2 into siRNAs, which are bound by AGO-2.

#### Exploratory analysis in Graphosoma lineatum

To assess whether the observed processing patterns are conserved across insect orders, an exploratory analysis was performed using RISC-bound small RNAs from the hemipteran species *Graphosoma lineatum* (Striped shield bug). Samples were collected 24 h after injection of 2 μg dsmGFP and analysed by strand-specific sequencing (Fig. 4).

**Figure 4.**
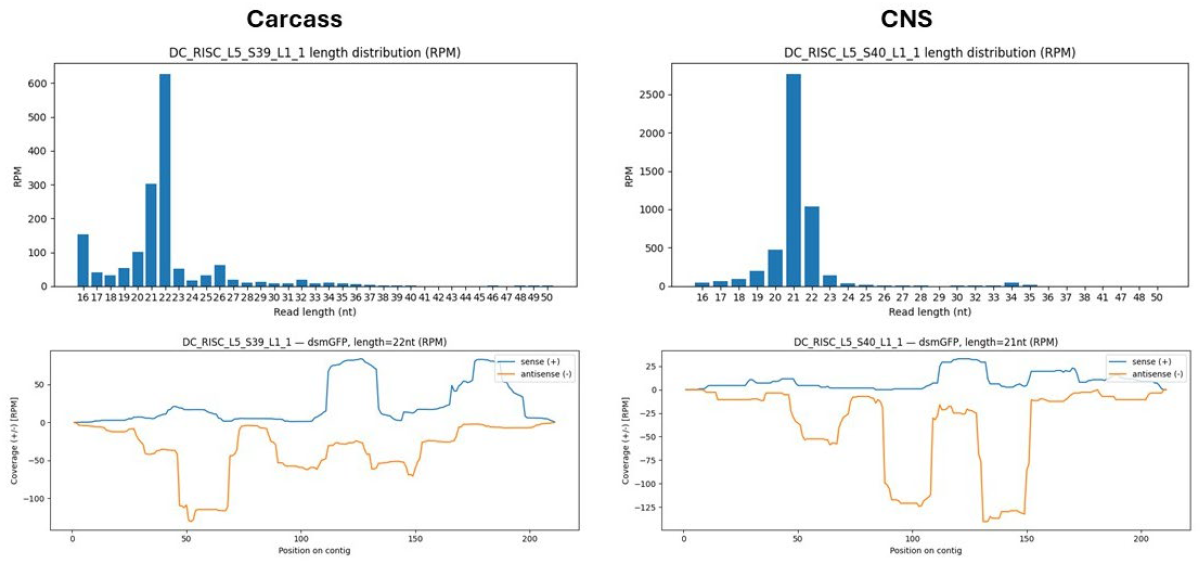
RISC-bound siRNA size distribution and mapping profiles in Graphosoma lineatum. (a) Length distribution of RISC-associated small RNAs recovered from the CNS 24 h after dsmGFP exposure, showing a dominant 21-nt siRNA peak. (b) Strand-specific mapping of 21-nt siRNAs from the CNS sample onto the dsmGFP template, revealing a major antisense hotspot similar to that observed in CPB tissues. (c) Length distribution of RISC-bound small RNAs from the remaining tissues, demonstrating a tissue-specific shift to a predominant 22-nt siRNA population. (d) Strand-specific mapping of 22-nt siRNAs from the remaining tissues sample, showing relocation of the dominant antisense hotspot to a secondary region of the dsmGFP sequence.

In CNS tissue (Fig. 4), a dominant population of 21-nt antisense siRNAs was detected, closely resembling the size distribution and mapping profile observed in CPB tissues (Fig. 2). These siRNAs mapped to the major hotspot region identified in CPB, indicating conserved processing preferences in neural tissue.

In contrast, remaining tissues samples displayed a shift towards a predominant 22-nt antisense siRNA population, accompanied by relocation of the dominant hotspot to a secondary processing region. This tissue-specific shift in both siRNA length and hotspot position suggests that Dicer activity or AGO loading preferences may differ between neural and peripheral tissues in *G. lineatum* and from those observed in CPB.

Although based on a single biological replicate (with a pool of 25 insects), these findings provide strong preliminary evidence that dsRNA-derived siRNA processing may vary across tissues and species. Further studies will be required to determine the mechanistic basis and biological significance of these differences.

## Conclusions

This study demonstrates that orally taken up dsRNA is predominantly processed into 21-nucleotide siRNAs across midgut, remaining tissues, and central nervous system tissues of *L. decemlineata*, indicating conserved Dicer-2-mediated processing throughout peripheral and neural compartments. Strand-specific mapping revealed discrete and stable cleavage hotspots along the dsmGFP sequence that were maintained across tissues, despite substantial differences in siRNA abundance. These findings indicate that dsRNA processing specificity is largely independent of tissue context and is primarily determined by intrinsic properties of the dsRNA substrate.

The detection of RISC-bound siRNAs in neural tissue provides molecular evidence for effective systemic delivery and canonical processing of orally administered dsRNA in the CNS. Although siRNA accumulation varied markedly among tissues, dsmGFP-derived sequences represented only a minor fraction of the total small RNA pool, suggesting that the RNAi machinery was not saturated under the experimental conditions. Exploratory analysis in *Graphosoma lineatum* further indicates that both conserved and tissue-specific features of dsRNA processing may occur across insect orders. Collectively, these results provide mechanistic insight into systemic RNAi in insects and support the potential of orally delivered dsRNA for functional and applied applications.

## Methods

### dsRNA synthesis

A template for the previously established control dsRNA sequence dsmGFP^26^ (Table S 1) was obtained from IDT (Germany) as an eBlock. Primer pair containing the T7 promoter sequence (“GAATTGTAATACGACTCACTATAGG”) at the 5 ʹ ends were designed to amplify dsmGFP. PCR reactions (50 μL) were performed using Q5® Hot Start High-Fidelity 2X Master Mix with an annealing temperature of 60°C for 30 cycles. Amplification yielded a single product, which was confirmed by agarose gel electrophoresis and purified using the Gel and PCR Clean-up kit (Macherey–Nagel, Germany).

dsRNA synthesis was carried out in 20 μL reactions using the MEGAscript™ T7 Transcription Kit (Invitrogen, Germany), followed by purification via lithium chloride precipitation^26^. Purified dsRNAs were resuspended in nuclease-free water, and concentrations were determined by Nanodrop spectroscopy (45 μg/OD260). RNA strands were annealed by heating to 94°C for 5 min and cooling at room temperature for 30 min. The resulting annealed dsRNAs (2.5 μg) were verified on a 1.5% agarose gel alongside a dsRNA ladder (NEB# N0363S, Germany).

### Insect Rearing and dsRNA Delivery

Adult Colorado potato beetles (*Leptinotarsa decemlineata*, CPB) were reared under controlled temperature and humidity conditions (25 °C, 65% humidity, 16 h light: 8 h dark) in greenhouse facilities. For CPB feeding experiments, thirty adult insects were starved overnight and subsequently placed individually on potato leaf disks (18 mm diameter) coated with 30 μL of 2 μg dsRNA targeting GFP (dsmGFP) solution containing 200 ppm Triton X-100. Feeding continued for 24 hours, after which midgut, brain (including CNS), and remaining tissues tissues were dissected from fifteen insects per biological replicate and pooled (two replicates per tissue type).

*Graphosoma lineatum* adults were collected from natural habitats in the Sillenbuch area of Stuttgart, Germany, in June 2025 and maintained in ventilated plastic containers for two days under controlled conditions (16:8 light cycle, 65% relative humidity) while being fed carrots and foliage of Apiaceae from their native habitat. For injection assays, thirty adult insects were injected with dsmGFP into the haemolymph using glass capillary needles in accordance with protocols established in Prof. Bucher’s laboratory (University of Göttingen, Germany), targeting the ventral fourth/fifth abdominal tergum. Following a 24-hour incubation, CNS and remaining tissues tissues from twenty-five insects were dissected in ice-cold PBS and pooled per tissue (one biological replicate per tissue type). See Fig. 5 for summarized experimental workflow.

**Figure 5.**
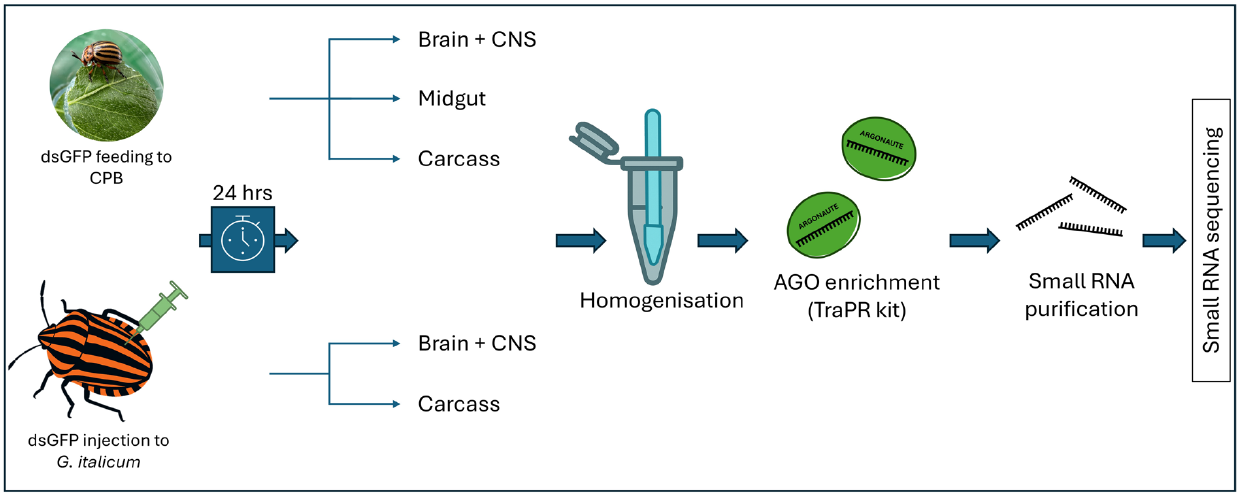
Illustration of study design and protocol. (note: AI generated images (G. italicum image and tubes) are used to create this illustration)

### Tissue Processing and RISC-bound Small RNA Isolation

Dissected tissues were immediately stored at –80 °C. Active RISC-bound small RNAs were purified using the TraPR Small RNA Isolation Kit (Lexogen), following the manufacturer’s instructions^25^. Samples were homogenized in lysis buffer using RNase-free micro-pestles, clarified by centrifugation, and applied to TraPR columns for isolation. RISC complexes were eluted, and small RNAs were extracted by phenol/chloroform precipitation and washed three times with ice-cold ethanol. Final RNA pellets were dissolved in 10 μL of elution buffer and stored at –80 °C until library preparation.

### Small RNA Library Preparation and Sequencing

Libraries from purified small RNAs (∼500 ng RNA per sample) were prepared using the NEBNext® Multiplex Small RNA Library Prep Kit (NEB #E7560S), following the manufacturer’s instructions. Final PCR amplification was performed with 11 cycles, and DNA products were purified using the Monarch PCR & DNA Clean-up Kit (NEB) and size-selected (135–150 bp cDNA) by 6% polyacrylamide gel electrophoresis, corresponding to 16–30 nt small RNAs. Library quality was assessed using an Agilent 2100 Bioanalyzer. Pooled libraries were sequenced at BGI using the DNBSEQ-G400 platform.

## Data Processing and Strand-Specific Mapping

Raw sequencing reads were trimmed and quality-filtered (Phred score > 30) using TrimGalore (v0.6.1, https://github.com/FelixKrueger/TrimGalore). Cleaned reads were mapped onto dsmGFP sequence using Bowtie 1 (v1.3) with zero-mismatch settings (–all, –n 0) to ensure high specificity. Mapped reads of 21-nt length were extracted and analysed using the samtools (v1.23) package for read count normalization and downstream tissue-specific small RNA profiling.

## Acknowledgement

The authors thank Dr. Georg Petschenka for adding valuable comments that improved the manuscript. The authors also thank Dr. Hamdow Al Karrat for assistance with collecting *Graphosoma lineatum* insects in the field. We further thank Dr. Gözde Güney for preparing the small RNA libraries. D.C. was supported by the Deutscher Akademischer Austauschdienst (DAAD) research grants program, which partially funded this work.

## Author Contributions

V.P.S.A. conceived and designed the study. V.P.S.A. performed insect experiments, tissue dissection, and small RNA isolation. D.C. processed small RNA for sequencing and conducted bioinformatic analyses. V.P.S.A. drafted the manuscript. Both authors contributed equally to data interpretation, revised the manuscript, and approved the final version.

## Declaration

During the preparation of this manuscript, the author (VPSA) used Perplexity tool to improve grammar, spelling, and vocabulary for clarity and readability. Perplexity was not used to generate new scientific content, results, analyses, or interpretations. The authors reviewed and edited all text as needed and takes full responsibility for the accuracy and integrity of the published work.

## References

1. Palli, S. R. RNAi turns 25:contributions and challenges in insect science. Front. Insect Sci. 3, 1209478 (2023).

2. Rank, A. P. & Koch, A. Lab-to-Field Transition of RNA Spray Applications – How Far Are We? Front. Plant Sci. 12, 755203 (2021).

3. Lau, P.-W. et al. The molecular architecture of human Dicer. Nat Struct Mol Biol 19, 436–440 (2012).

4. Hammond, S. M., Boettcher, S., Caudy, A. A., Kobayashi, R. & Hannon, G. J. Argonaute2, a Link Between Genetic and Biochemical Analyses of RNAi. Science 293, 1146–1150 (2001).

5. Filipowicz, W., Bhattacharyya, S. N. & Sonenberg, N. Mechanisms of post-transcriptional regulation by microRNAs: are the answers in sight? Nat Rev Genet 9, 102–114 (2008).

6. Neumeier, J. & Meister, G. siRNA Specificity: RNAi Mechanisms and Strategies to Reduce Off-Target Effects. Front. Plant Sci. 11, 526455 (2021).

7. De Neef, E. et al. A bioinformatic ecological risk assessment framework for externally applied double-stranded RNA-based biopesticides. Integrated Environmental Assessment and Management 22, 116–131 (2026).

8. Tardin-Coelho, R. et al. A systematic review on public perceptions of RNAi-based biopesticides: Developing Social Licence to Operate. npj Sustain. Agric. 3, 15 (2025).

9. Pallis, S. et al. Effects of Low Doses of a Novel dsRNA-based Biopesticide (Calantha) on the Colorado Potato Beetle. Journal of Economic Entomology 116, 456–461 (2023).

10. Rodrigues, T. B. et al. First Sprayable Double-Stranded RNA-Based Biopesticide Product Targets Proteasome Subunit Beta Type-5 in Colorado Potato Beetle (Leptinotarsa decemlineata). Front. Plant Sci. 12, 728652 (2021).

11. Palli, S. R. RNA interference in Colorado potato beetle: steps toward development of dsRNA as a commercial insecticide. Current Opinion in Insect Science 6, 1–8 (2014).

12. Mishra, S., Lamour, K., Emrich, S., Moar, W. & Jurat-Fuentes, J. L. Reduced uptake through clathrin down-regulation is associated with resistance to dsRNA in a population of the Colorado potato beetle (Leptinotarsa decemlineata, Say). Pesticide Biochemistry and Physiology 216, 106783 (2026).

13. Mehlhorn, S. G., Geibel, S., Bucher, G. & Nauen, R. Profiling of RNAi sensitivity after foliar dsRNA exposure in different European populations of Colorado potato beetle reveals a robust response with minor variability. Pesticide Biochemistry and Physiology 166, 104569 (2020).

14. Petek, M., Coll, A., Ferenc, R., Razinger, J. & Gruden, K. Validating the Potential of Double-Stranded RNA Targeting Colorado Potato Beetle Mesh Gene in Laboratory and Field Trials. Front. Plant Sci. 11, 1250 (2020).

15. Cappelle, K., De Oliveira, C. F. R., Van Eynde, B., Christiaens, O. & Smagghe, G. The involvement of clathrin-mediated endocytosis and two Sid-1-like transmembrane proteins in double-stranded RNA uptake in the Colorado potato beetle midgut. Insect Molecular Biology 25, 315–323 (2016).

16. Shi, X. et al. Cellular uptake of extracellular dsRNA is tissue-dependent in insects. BMC Biol https://doi.org/10.1186/s12915-026-02526-6 (2026) doi:10.1186/s12915-026-02526-6.

17. Wynant, N., Verlinden, H., Breugelmans, B., Simonet, G. & Vanden Broeck, J. Tissue-dependence and sensitivity of the systemic RNA interference response in the desert locust, Schistocerca gregaria. Insect Biochemistry and Molecular Biology 42, 911–917 (2012).

18. Limmer, S., Weiler, A., Volkenhoff, A., Babatz, F. & KlÃ¤mbt, C. The Drosophila blood-brain barrier: development and function of a glial endothelium. Front. Neurosci. 8, (2014).

19. Schirmeier, S. & Klämbt, C. The Drosophila blood-brain barrier as interface between neurons and hemolymph. Mechanisms of Development 138, 50–55 (2015).

20. Carlson, S. D., Juang, J.-L., Hilgers, S. L. & Garment, M. B. Blood Barriers of the Insect. Annu. Rev. Entomol. 45, 151–174 (2000).

21. Hindle, S. J. & Bainton, R. J. Barrier mechanisms in the Drosophila blood-brain barrier. Front. Neurosci. 8, (2014).

22. Groen, S. C. et al. Multidrug transporters and organic anion transporting polypeptides protect insects against the toxic effects of cardenolides. Insect Biochemistry and Molecular Biology 81, 51–61 (2017).

23. Petschenka, G., Pick, C., Wagschal, V. & Dobler, S. Functional evidence for physiological mechanisms to circumvent neurotoxicity of cardenolides in an adapted and a non-adapted hawk-moth species. Proc. R. Soc. B. 280, 20123089 (2013).

24. Hernández-Pelegrín, L., Mishra, S., Jurat-Fuentes, J. L. & Herrero, S. Antiviral RNAi is preserved in dsRNA-resistant Colorado potato beetle (Leptinotarsa decemlineata). Insect Biochemistry and Molecular Biology 187, 104469 (2026).

25. Grentzinger, T. et al. A universal method for the rapid isolation of all known classes of functional silencing small RNAs. Nucleic Acids Research 48, e79–e79 (2020).

26. Cedden, D., Güney, G., Rostás, M. & Bucher, G. Optimizing dsRNA sequences for RNAi in pest control and research with the dsRIP web platform. BMC Biol 23, 114 (2025).

27. Guan, R., Hu, S., Li, H., Shi, Z. & Miao, X. The in vivo dsRNA Cleavage Has Sequence Preference in Insects. Front. Physiol. 9, 1768 (2018).

