## Supplementary material for "Orally Delivered dsRNA-Derived siRNAs Reach the Central Nervous System in *Leptinotarsa decemlineata*": Table S 1, Table S 2

| | |
|---|---|
| dsmGFP | GGACCCTGACCTACGGCCTATTAGTGTGATCGATACTACCACTGACGTACAATG AGATCACGAATCAAGAAGATAGTGTCAGCACCAGGAGCGCACCATCTTAGTCA TCAGCAGACGCATCGACTCTAGACAGCTACGCCGAGGTGAAGTTCGAGCGACA CCCTGGTGAACCGCAATACTGTAGAGCATCAACTAGTACGAGGACGGCACC |
| dsmGFP_F_T7 | GAATTGTAATACGACTCACTATAGGTGCCGTCCTCGTACTAGTT |
| dsmGFP_R_T7 | GAATTGTAATACGACTCACTATAGGACCCTGACCTACGGCCTAT |

*Table S 1 dsRNA target sequence and primers used for dsmGFP synthesis. The full nucleotide sequence of the dsmGFP fragment used to generate dsRNA* used in this study *is shown. Forward and reverse primers containing the T7 promoter sequence (5'-TAATACGACTCACTATAGG-3') were used to amplify the dsmGFP template for in vitro transcription.*

| total_reads_for_norm | mapped_total_raw | Proportion of dsmGFP per total reads | Tissue_sample | sample | dsmGFP_normalisation |
|---|---|---|---|---|---|
| 21798257 | 12000 | 0.055050273 | Carcass_CPB | DC_RISC_L5_S33_L1_1 | 0.000550503 |
| 17448409 | 8741 | 0.050096258 | Carcass_CPB | DC_RISC_L5_S34_L1_1 | 0.000500963 |
| 4451394 | 46621 | 1.047334835 | Midgut_CPB | DC_RISC_L5_S35_L1_1 | 0.010473348 |
| 9528383 | 159873 | 1.677860766 | Midgut_CPB | DC_RISC_L5_S36_L1_1 | 0.016778608 |
| 3642390 | 4758 | 0.130628516 | CNS+BRAIN_CPB | DC_RISC_L5_S37_L1_1 | 0.001306285 |
| 3673534 | 3600 | 0.097998276 | CNS+BRAIN_CPB | DC_RISC_L5_S38_L1_1 | 0.000979983 |
| 21092310 | 33711 | 0.159826022 | Carcass_stinkbug | DC_RISC_L5_S39_L1_1 | 0.00159826 |
| 1124677 | 5597 | 0.497653993 | CNS+BRAIN_stinkbug | DC_RISC_L5_S40_L1_1 | 0.00497654 |

*Table S 2 Summary of AGO-enriched small RNA sequencing and dsmGFP-derived read counts.For each tissue sample, the table reports total reads used for normalization, the number of raw reads mapping to the dsmGFP sequence, and the proportion of dsmGFP-mapping reads relative to total reads. The normalized dsmGFP abundance (dsmGFP_normalisation) is provided to enable comparisons across samples and tissues.*
